# CelloType: A Unified Model for Segmentation and Classification of Tissue Images

**DOI:** 10.1101/2024.09.15.613139

**Authors:** Minxing Pang, Tarun Kanti Roy, Xiaodong Wu, Kai Tan

## Abstract

Cell segmentation and classification are critical tasks in spatial omics data analysis. We introduce CelloType, an end-to-end model designed for cell segmentation and classification of biomedical microscopy images. Unlike the traditional two-stage approach of segmentation followed by classification, CelloType adopts a multi-task learning approach that connects the segmentation and classification tasks and simultaneously boost the performance of both tasks. CelloType leverages Transformer-based deep learning techniques for enhanced accuracy of object detection, segmentation, and classification. It outperforms existing segmentation methods using ground-truths from public databases. In terms of classification, CelloType outperforms a baseline model comprised of state-of-the-art methods for individual tasks. Using multiplexed tissue images, we further demonstrate the utility of CelloType for multi-scale segmentation and classification of both cellular and non-cellular elements in a tissue. The enhanced accuracy and multi-task-learning ability of CelloType facilitate automated annotation of rapidly growing spatial omics data.

## Introduction

Recent advancements in spatial omics technologies have markedly improved our ability to analyze intact tissues at the cellular level, revealing unparalleled insights into the link between cellular architecture and functionality of various tissues and organs^1^. Collaborative efforts, such as the Human Tumor Atlas Network^2^, the Human Biomolecular Atlas Program^3^, and the BRAIN initiative, are leveraging these technologies to map spatial organizations of various types of healthy and diseased tissues. With the anticipated surge in spatial omics data, there is a pressing need for sophisticated computational tools for data analysis. A typical analysis workflow of spatial omics data begins with cell segmentation. Following cell segmentation and quantification of molecular analytes, cell type annotation is the next critical, albeit often time-consuming task before further analysis can proceed. Conventional analysis pipelines perform these two tasks sequentially, typically using the segmentation results as the inputs for the classification task. As representatives of state-of-the-art segmentation methods, Mesmer^4^ uses a convolutional neural network (CNN)^5^ backbone and a Feature Pyramid Network with the watershed algorithm for both nuclear and cell segmentation. Cellpose^6^ and Cellpose2^7^ use a CNN with a U-net^8^ architecture to predict the gradient of topological map. A gradient tracking algorithm is then used to obtain the segmentation mask. For cell classification task, CellSighter^9^ employs CNN to predict cell types based on segmentation masks and the tissue images. CELESTA^10^ uses an iterative algorithm to assign cell types based on quantified cell-by-protein matrix.

Despite achieving satisfactory performance in certain tissues, conventional approaches have several limitations. First and foremost, the reliance of cell classification models on segmentation results hampers their ability to leverage the full spectrum of semantic information present in tissue images. In fact, these two tasks are interconnected. Segmentation can enhance focus on relevant signals, thus mitigating noise and enabling more precise learning of class features for classification. Conversely, information specific to classes aid in the segmentation process, as the unique texture and morphology of certain object types can enhance segmentation accuracy.

Second, the two-step approach is computationally inefficient, requiring separate training for each task. Third, the performance of existing segmentation methods also varies significantly across different tissue types, suggesting substantial room for improvement. Moreover, to our knowledge, existing methods do not offer a confidence assessment for the segmentation task.

Deep learning, especially through the use of CNNs, has gained popularity in biomedical image analysis, especially in segmentation^11^ and classification^9^. Mesmer, for example, has notably improved cell segmentation accuracy using CNN. However, recent developments in computer vision has shown that Transformer-based models^12^, such as the Detection Transformer (DETR) ^13^ and the Detection transformer with Improved deNoising anchOr (DINO)^14^, significantly outperform CNN-based models in object detection. These Transformer-based models have also shown superior performance in instance segmentation of histological images^15^. Despite these breakthroughs, the application of Transformer-based models to cell/nuclear segmentation in multiplexed images and other spatial omics data type remains unexplored. A unified framework, MaskDINO^16^, which integrates object detection and segmentation, has shown superior performance across diverse datasets for multi-class instance segmentation. However, its effective ness has only been tested on RGB images of natural objects. This leaves a significant gap in applying Transformer-based models to multiplexed tissue images, which present greater challenges due to their larger number of imaging channels, varying shapes of tightly apposed/overlapping cellular and non-cellular elements.

The limitations of current methodologies and the advent of novel deep learning techniques motivated us to develop CelloType, an end-to-end method for joint cell segmentation and classification. CelloType employs a Transformer-based deep neural network architecture with multiple branches to handle object detection, segmentation, and classification concurrently. We benchmarked the performance of CelloType against state-of-the-art methods using a variety of public image datasets, including single-channel, and multiplexed fluorescent tissue and cell images and bright-field images of nature objects. We further demonstrated a novel feature of CelloType for multi-scale segmentation and classification to delineate both cellular and noncelluar elements in tissue images.

## Results

### Overview of CelloType

CelloType is a deep neural network (DNN)-based framework (Figure 1) designed for joint multi- scale segmentation and classification of a variety of biomedical microscopy images, including multiplexed molecular images, histological images, and bright-field images. The core of CelloType’s functionality begins with the extraction of multi-scale image features through the use of a Swin Transformer^17^. These features are then fed into the DINO object detection module that extracts instance-specific latent features and predicts a preliminary object bounding box with associated class label for each instance. Finally, the MaskDINO segmentation module integrates the multi-scale image features from the Swin Transformer and DINO outputs to produce the final refined instance segmentations. The CelloType model is trained using a loss function that considers segmentation masks, object detection boxes, and classes labels.

**Figure 1 –.**
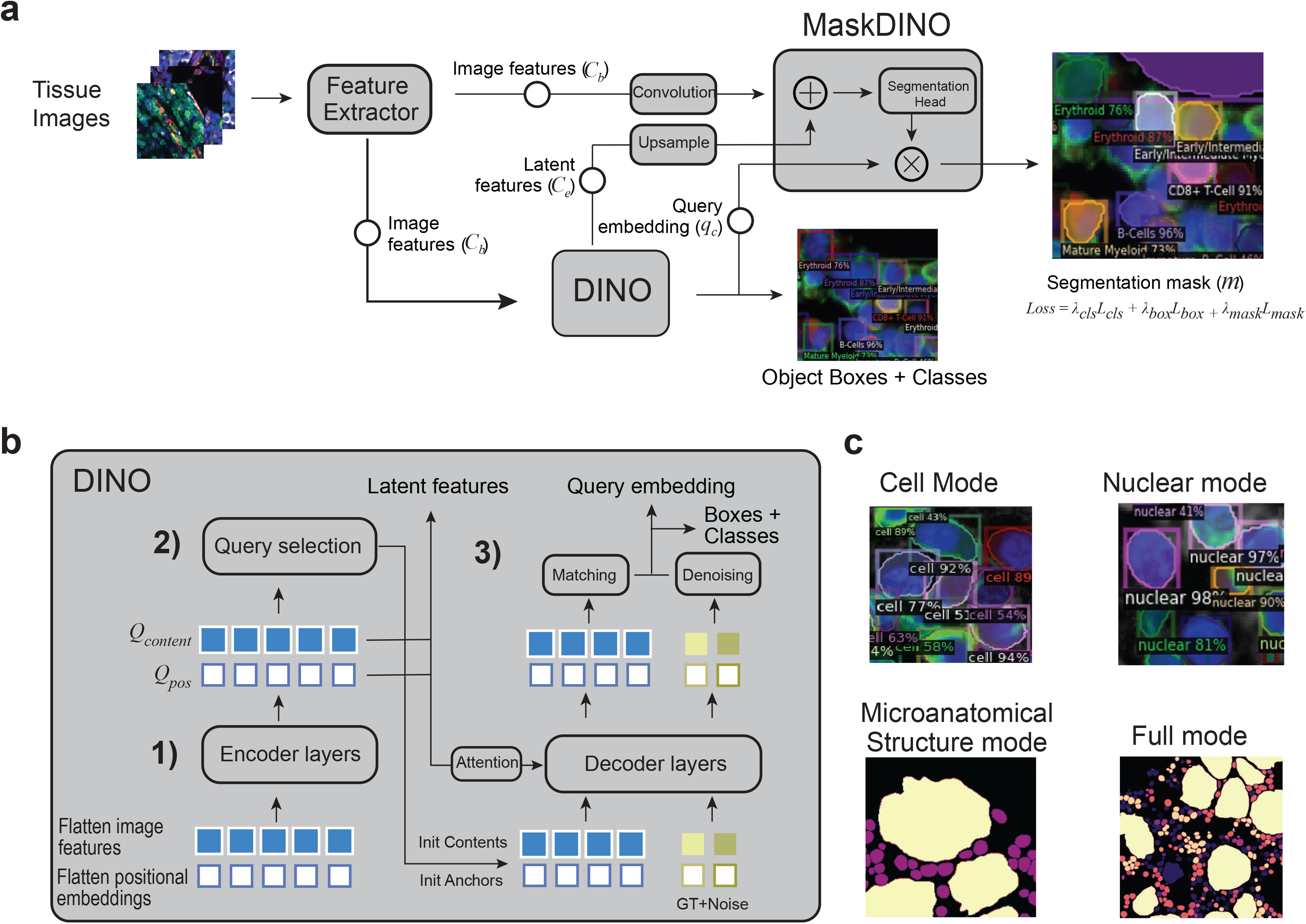
Overview of CelloType. **a)** Overall architecture, input, and output of CelloType. First, a Transformer-based feature extractor is employed to derive multi-scale features (*C_b_*) from the image. Second, using a Transformer-based architecture, the DINO object detection module extracts latent features (*C_e_*) and query embeddings (*q_c_*) that are combined to generate object detection boxes with cell type labels. Subsequently, the MaskDINO module integrates the extracted image features with DINO’s outputs, resulting in detailed instance segmentation and cell type classification. During training, the model is optimized based on an overall loss function (*Loss*) that considers losses based on cell segmentation mask (*λ_mask_L_mask_*), bounding box (*λ_box_L_box_*), and cell type label (*λ_cls_L_cls_*). **b**) Input, output, and architecture of the DINO module. The DINO module consists of a multi-layer Transformer and multiple prediction heads. DINO starts by flattening the multi- scale features from the Transformer-based feature extractor. These features are merged with positional embeddings to preserve spatial context (step 1 in the figure). DINO then employs a mixed query selection strategy, initializing positional queries (*Q_pos_*) as anchor detection boxes and maintaining content queries (*Q_content_*)as learnable features, thus adapting to the diverse characteristics of cells (step 2). The model refines these anchor boxes through decoder layers using deformable attention mechanism and employs contrastive denoising training by introducing noise to ground truth (GT) labels and boxes to improve robustness and accuracy. Then a linear projection acts as the classification branch to produce the classification results for each box (step 3). **c**) Multi-scale ability of CelloType. CelloType is versatile and can perform a range of end-to-end tasks at different scales, including cell segmentation, nuclear segmentation, microanatomical structure segmentation, and full instance segmentation with corresponding class annotations.

The DINO module’s architecture (Figure 1b) includes a Transformer encoder-decoder set-up with multiple prediction heads. It begins by flattening image features and integrating them with positional embeddings^18^. By employing a strategy that mixes anchor and content queries, the module can adapt to various object features. The module refines bounding boxes through a deformable attention mechanism. A contrastive denoising training (CDN) procedure is used together with the attention mechanism to improve the robustness of bounding box detection.

Finally, a linear transformation is applied to the denoised bounding box features to predict the class label of the object.

CelloType can tackle diverse image analysis tasks including cell/nuclear segmentation, non-cellular structure segmentation, and multi-scale segmentation (Figure 1c). Different data types are used to train CelloType for various tasks. For cell or nuclear segmentation, training data includes one/two-channel images with corresponding cell membrane or nuclear masks. For joint segmentation and classification, the training data consists of images with segmentation mask, bounding box, and class label of each object. The images can contain many channels in addition to the cell membrane and nuclear channels. CelloType is implemented in Python and publicly available at http://github.com/tanlabcode/CelloType.

### Benchmark of cell and nuclear segmentation performance using multiplexed images

We first applied CelloType to the TissueNet dataset^4^ that includes tissue images generated using six multiplexed molecular imaging technologies (CODetection by indexing (CODEX)^19^, Cyclic Immunofluorescence (CycIF)^20^, Imaging Mass Cytometry (IMC)^21^, Multiplexed Ion Beam Imaging (MIBI)^22^, Multiplexed Immunofluorescence (MxIF)^23^, and Vectra^24^) and six tissue types (breast, gastrointestinal, immune, lung, pancreas, skin). The images were divided into 2,580 training patches (512 x 512 pixels) and 1,324 test patches (256 x 256 pixels).

We compared CelloType with two state-of-the-art methods, Mesmer^4^ and Cellpose2^7^. For object detection and instance segmentation, we used the Average Precision (AP) metric^25^ defined by the Common Objects in Context (COCO) project and the Intersection over Union (IoU) thresholds from 0.5 to 0.9 in 0.05 increments (Methods). The precision-IoU curves (Figure 2a) revealed that CelloType consistently outperformed both Mesmer and Cellpose2 across the entire range of IoU thresholds on the TissueNet dataset. Additionally, considering that CelloType provides a confidence score for each segmentation mask and the COCO metric incorporates these confidence scores in matching predicted and ground truth cell boundaries, we also evaluated a version of CelloType that outputs confidence scores, CelloType_C. Overall, performance is higher for cell segmentation than nuclear segmentation for all methods except for Mesmer. For cell segmentation, CelloType_C achieved an average AP of 0.556, significantly surpassing the basic CelloType (0.450), Cellpose2 (0.354), and Mesmer (0.312). For nuclear segmentation, CelloType_C achieved a mean AP of 0.655, outperforming CelloType (0.571), Cellpose2 (0.516), and Mesmer (0.237) by considerable margins. These results underscore CelloType’s superior segmentation accuracy and the added value of incorporating confidence scores.

**Figure 2 –.**
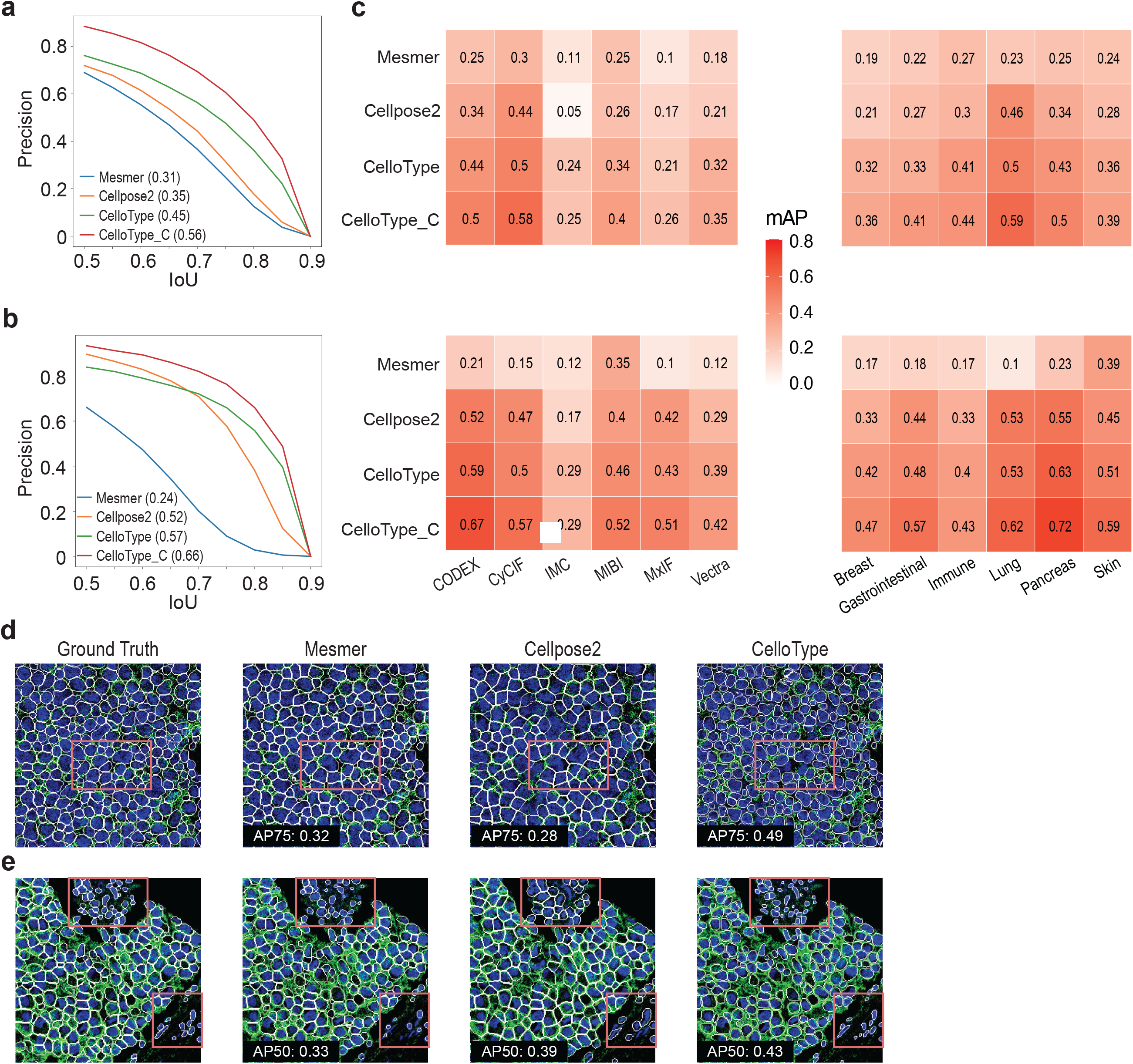
Evaluation of segmentation accuracy using TissueNet datasets. **a**) Average Precision (AP) across Intersection over Union (IoU) thresholds for cell segmentation by Mesmer, Cellpose2, CelloType and CelloType_C (CelloType with confidence score). Mean AP value across IoU thresholds of 0.5-0.9 (mAP) for each method is indicated in parenthesis. **b**) AP across IoU thresholds for nuclear segmentation. **c**) Performance of methods stratified by imaging platform and tissue type. The top left heatmap shows the mAP scores for cell segmentation stratified by imaging platform, including CODEX, CyCIF, IMC, MIBI, MxIF and Vertra. The top right heatmap shows the mAP scores for cell segmentation stratified by tissue type, including breast, gastrointestinal, immune, pancreas and skin. The second row of heatmaps shows the mAP values for nuclear segmentation. **d**) Representative examples of cell segmentation of immune tissue imaged using Vectra platform. Blue, nuclear channel; green, membrane channel; white, cell boundary. The red box highlights a representative region that the methods perform differently. The AP75 score (Average precision at IoU threshold of 0.75) is displayed on the images. **e**) Representative examples of nuclear segmentation of gastrointestinal tissue using the IMC platform. The AP50 scores are shown on the images.

To evaluate the effect of imaging technology and tissue type on the segmentation performance, we next analyzed the mean AP scores stratified by these two factors (Figure 2b). Overall, performance of all methods is lowest on the IMC data and breast tissue data. CelloType and CelloType_C consistently outperformed Mesmer and Cellpose2 across the technology platforms and tissue types. Figure 2d-e show representative cell and nuclear segmentation results by the compared methods. These examples illustrate Cellpose2 tends to produce segmentation boundaries that are larger than the ground truth and thus often under-segmentation. On the other hand, Mesmer tends to miss more cells or nuclei.

### Benchmark of cell segmentation performance using diverse image types

To further evaluate CelloType’s performance of cell segmentation across diverse microscopy images beyond multiplexed fluorescent images, we applied CelloType to the Cellpose Cyto dataset^6^ which include fluorescent, bright-field microscopy images of cells and images of natural objects. Since most of the images in this dataset contain only one channel and Mesmer was trained on two-channel image data, we only benchmarked the performance of CelloType, CelloType_C, and Cellpose2.

Across the entire dataset, CelloType_C achieved an average AP of 0.469, surpassing the performance of both CelloType (0.368) and Cellpose2 (0.322). This superiority is consistently observed across 6 diverse image sets (Figure 3b). Figure 3c shows representative segmentation results by Cellpose2 and CelloType for a single-channel image from the “Other microscopy” category. Consistent with the findings in Figure 2d with multiplexed IMC image, Cellpose2 exhibited a tendency for under-segmentation, while CelloType produced more precise segmentation boundaries. Additionally, Figure 3d shows the segmentation result for another single-channel image from the “Non-fluorescent” cell category, where CelloType demonstrated enhanced accuracy in both identifying the correct number of cells and delineating their boundaries, in contrast to Cellpose2, which tended to over-segment.

**Figure 3 –.**
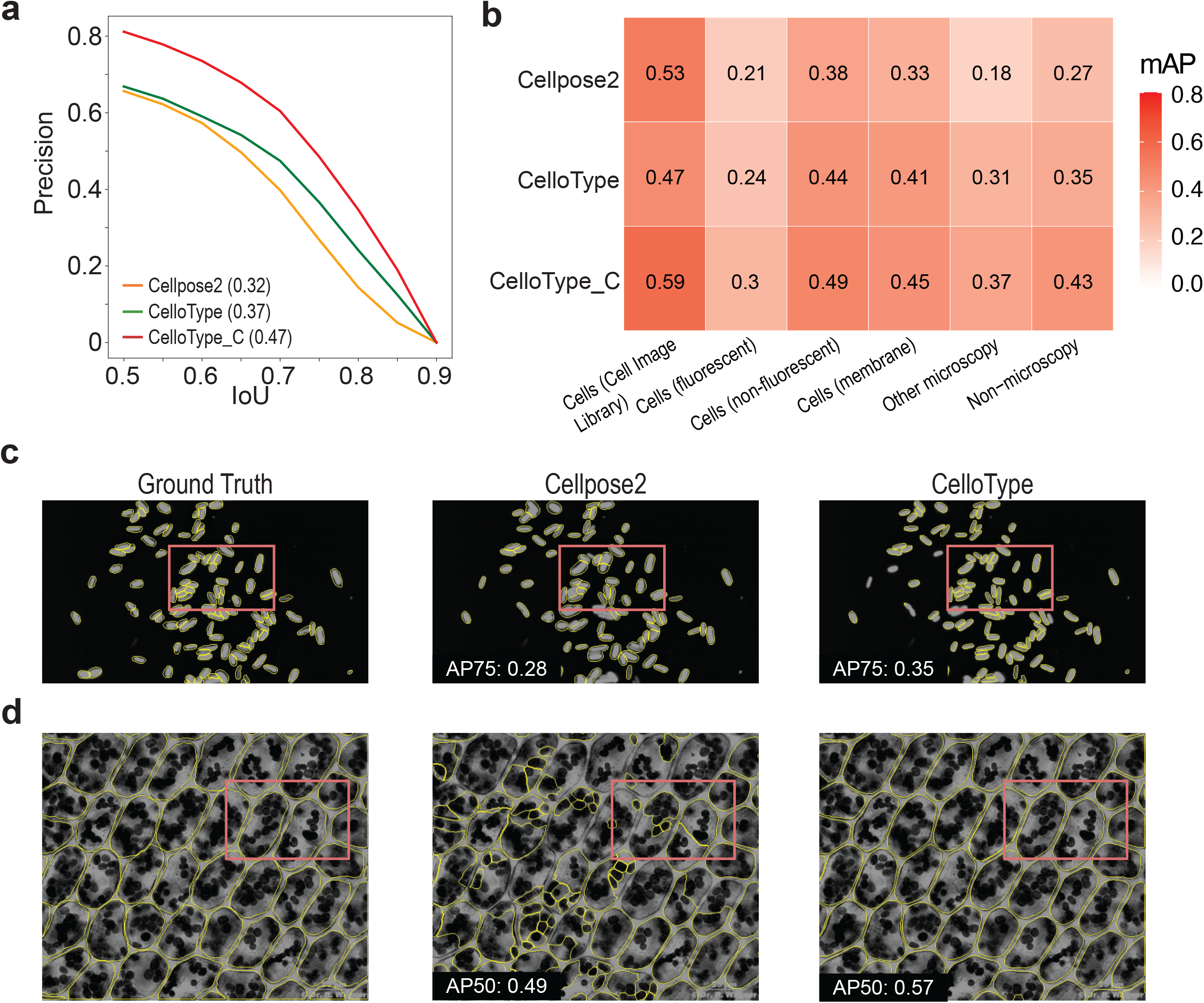
Evaluation of segmentation accuracy using Cellpose Cyto dataset. **a)** Average precision (AP) across Intersection over Union (IoU) thresholds for Cellpose2, CelloType and CelloType_C (CelloType with confidence score). Mean AP value across IoU thresholds of 0.5-0.9 (mAP) for each method is indicated in parenthesis. **b**) Mean AP values of Cellpose2, CelloType, and CelloType_C stratified by imaging modalities and cell types. The test dataset comprises microscopy and non-microscopy images from the Cellpose Cyto dataset that comprises 6 subsets, including Cells (Cell Image Library), Cells (Fluorecent), Cells (Non- fluorecent), Cells (Membrane), Other microscopy, and Non-microscopy. **c**) Representative examples of cell segmentation of a microscopy image by the compared methods. The red boxes highlight a representative region that the methods perform differently. The AP75 score is displayed on the images. **d**) Representative examples of cell segmentation of a non-fluorescent image by the compared methods.

### Joint segmentation and cell type classification of multiplexed images

To assess the performance of CelloType for simultaneous cell segmentation and classification, we applied it to a colorectal cancer CODEX dataset^26^. This dataset consists of 140 images of tumor tissue sections from 35 patients. Each tissue section was imaged using 56 fluorescent antibodies plus two nuclear stains, resulting in a total of 58 channels. These images were processed into 512 x 512 pixels image patches, which were subsequently divided into a training set of 720 patches and a test set of 120 patches (Supplemental Figure 1). Given the lack of established methods for simultaneous cell segmentation and classification, we combined Cellpose2 and CellSighter as a baseline model. This choice was motivated by the reported superior performance of each method for their respective task.

Using manual cell type annotation as the ground truth, we computed the AP score at an IoU threshold of 0.5 (i.e. AP50) for each cell type. CelloType achieved a mean AP50 of 0.84 across all cell types, markedly exceeding the Cellpose2+CellSighter model’s mean AP of 0.24 (Figure 4a). Furthermore, both CelloType_C and CellSighter produce a confidence score for their cell type predictions. To assess the utility of the confidence score, we explored the relationship between these confidence scores and accuracy of predictions. Notably, CelloType’s confidence scores demonstrated a strong, nearly linear correlation with prediction accuracy, particularly within the confidence score range of 0.5 to 0.7. In contrast, the relationship for CellSighter’s confidence scores appeared flat, indicating a lack of reliable calibration in its confidence assessment (Figure 4b).

**Figure 4 –.**
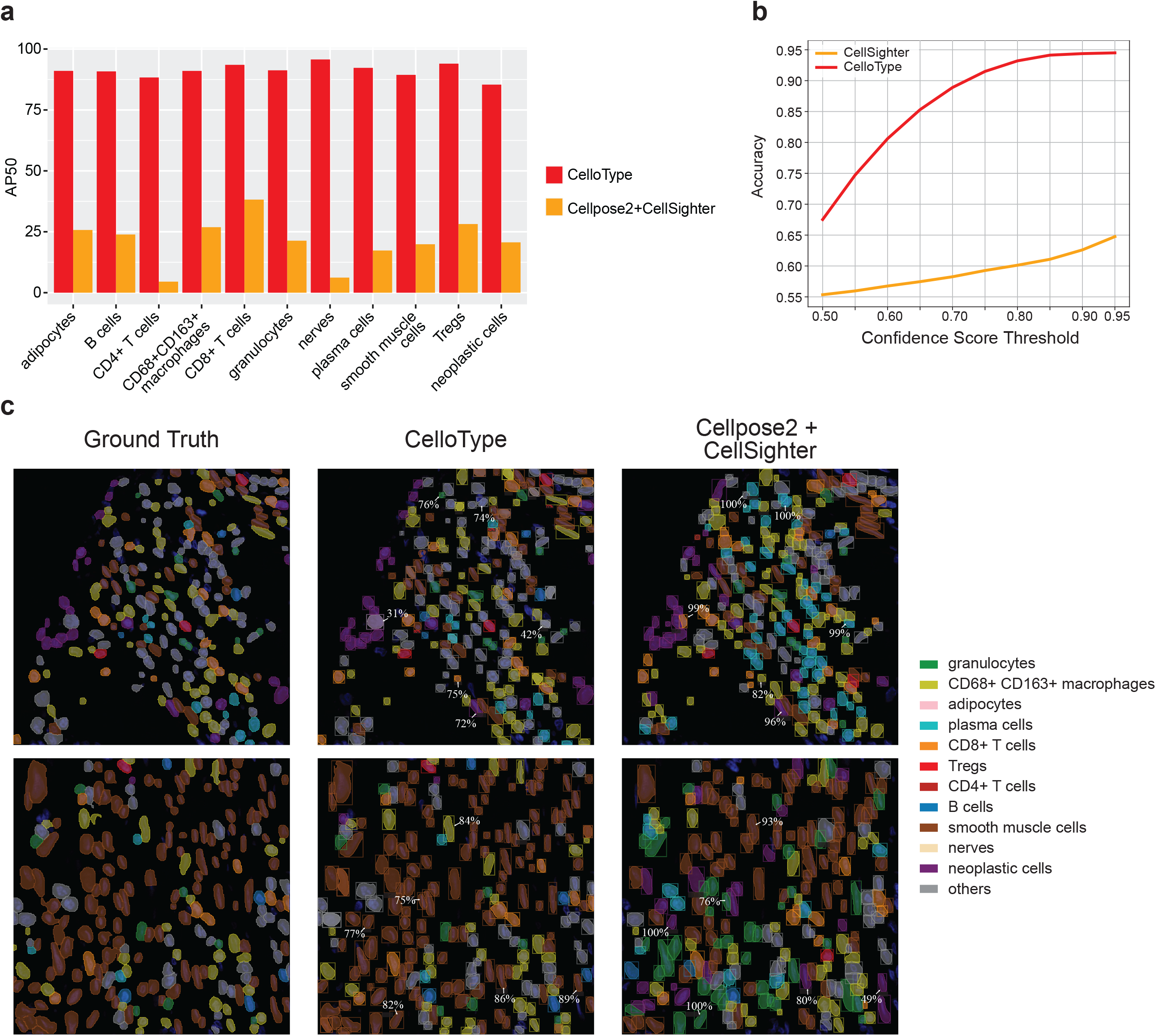
CelloType performs joint segmentation and cell type classification. **a)** Barplot showing AP50 values for cell type annotation by the two compared methods. **b)** Line plot showing the relationship between classification accuracy and confidence score threshold by the two methods. **c**) Representative examples of cell segmentation and classification results using the colorectal cancer CODEX dataset. Each row represents a 200 by 200 pixels field of view (FOV) of a CODEX image. Each FOV shows predicted cell segmentation masks (boxes) and cell types (colors). Ground Truth, manually annotated cell types; CelloType, end-to-end cell segmentation and cell type classification; Cellpose2+CellSighter, cell segmentation by Cellpose 2 followed by cell type classification by CellSighter. Randomly selected confidence scores for cell classification computed by the two methods were displayed next to the predicted instances.

Figure 4c shows two examples of predictions by CelloType and Cellpose2+CellSighter along with the ground truth annotations. These predictions encompass cell segmentation masks, predicted cell types and associated confidence scores. CelloType correctly predicted the identities of the vast majority of cells of different types with varying morphologies and abundance. For instance, in the top image, CelloType correctly predicted abundant neoplastic cells, alongside rare regulatory T cells (Treg), and morphologically irregular macrophages.

Similarly, in the bottom image, CelloType correctly predicted abundant smooth muscle cells and sparsely distributed CD8+ T cells. In contrast, the Cellpose2+CellSighter model misclassified several cell types as plasma cells (top image) and granulocytes (bottom image). Moreover, we found many instances where CellSighter’s predictions, despite being incorrect, were accompanied by high confidence scores, as indicated by arrows.

We next evaluated the performance of each component of the Cellpose2+CellSigher model, focusing on the segmentation function of Cellpose2 and the cell type classification function of CellSighter. Figure 5a shows the AP-IoU curve for cell segmentation on the colorectal cancer CODEX dataset. CelloType achieved a mean AP of 0.585, significantly exceeding Cellpose2’s mean AP of 0.345. In assessing CellSighter’s classification performance, we used the ground truth segmentation masks as inputs, treating the task purely a classification task. The resulting confusion matrix revealed the distribution of predictions for each cell type and the accuracy values displayed along the diagonal (Figure 5b). Furthermore, Figure 5c shows CellSighter’s classification precision for 11 cell types, achieving a mean precision of 0.53, compared to CelloType’s mean AP50 score of 0.81. This comparative analysis underscores CelloType’s superior performance not only as an end-to-end tool for cell type annotation but also in its individual functions for segmentation and classification, outperforming the two-stage approach of combining Cellpose2 and CellSighter.

**Figure 5 –.**
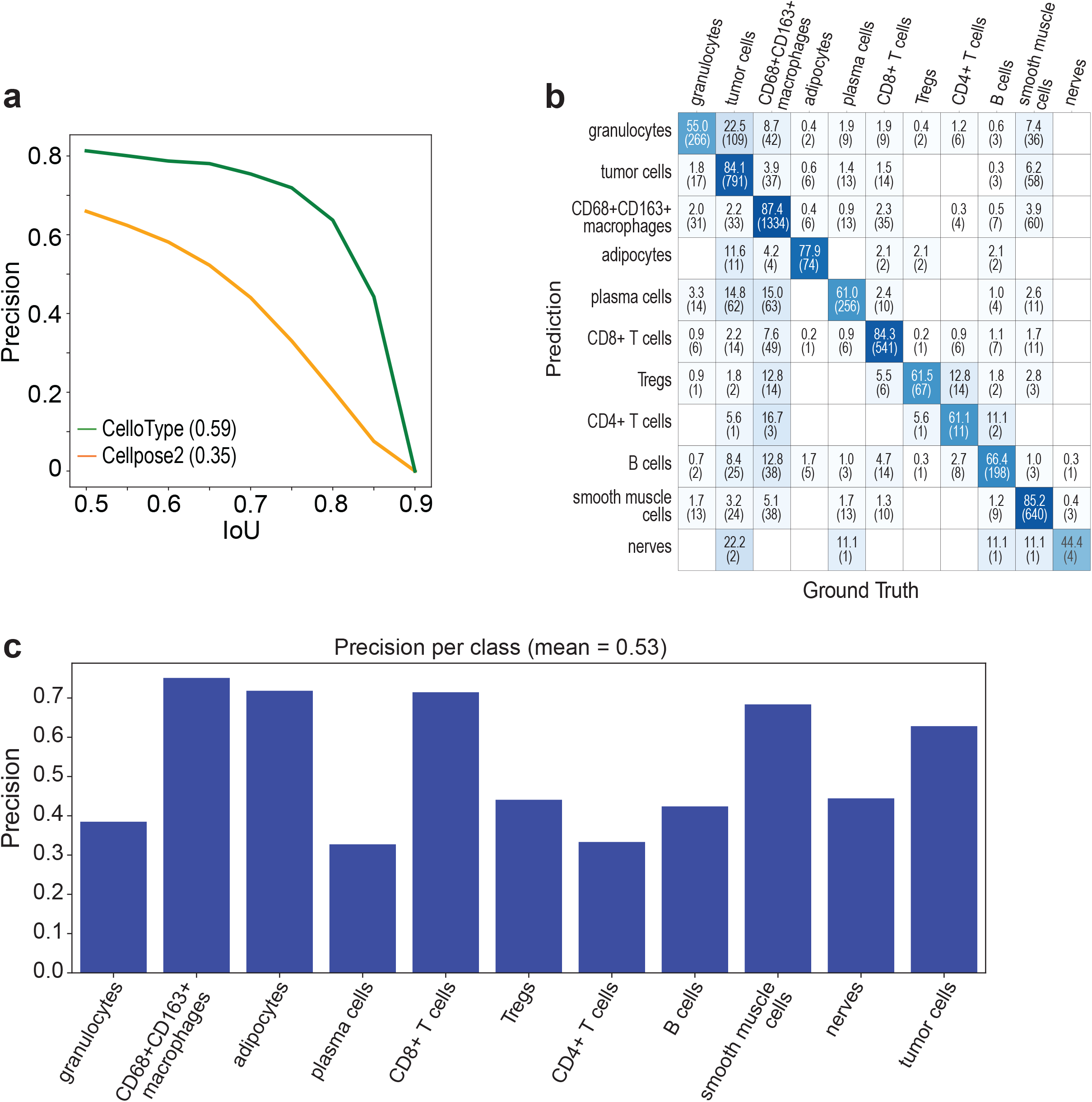
Performance benchmarking of Cellpose2 and CellSighter. Each method was evaluated for its originally intended task, namely Cellpose2 for segmentation and CellSighter for cell classification. Colorectal cancer CODEX dataset was used for benchmarking purpose. **a**) AP value of segmentation across a range of IoU thresholds. Mean AP value (mAP) is shown in parenthesis. **b)** Heatmap showing the confusion matrix of CellSighter cell type classification results. Ground truth cell segmentation masks were used as input to CellSighter. Each grid in the heatmap includes an accuracy score and the count of cells. **c)** Barplot showing the precision scores for each class identified by the CellSighter model based on the ground truth cell segmentation mask, with an overall mean precision of 0.53.

### Multi-scale segmentation and annotation by CelloType

Non-cellular components, such as the vasculature, lymphatic vessels, trabecular bone, and extra cellular matrix, and reticular fibers play important roles in tissue function. These elements are typically much larger than cells. Moreover, certain cell types like macrophages and adipocytes are either large or possess irregular shapes. Together, these elements present challenges to conventional segmentation methods. Furthermore, existing methods are incapable of simultaneous, multi-scale segmentation of both cellular and non-cellular elements within a tissue image. To assess the effectiveness of CelloType for multi-scale segmentation and classification, we applied it to a human bone marrow CODEX dataset^27^ (Supplemental Figure 2). This dataset comprises 12 whole-slide images of bone marrow sections from healthy donors, with each tissue section imaged using 53 fluorescent antibodies plus one nuclear stain, totaling 54 channels. The images were divided into 512 x 512 pixels patches with 1600 for training and 400 for testing.

The dataset presents a unique challenge due to the diversity of cell/non-cell types, notably adipocytes, which are substantially larger than other cell types, and trabecular bone fragments, which have irregular and complex shapes.

Using 5-fold cross-validation, we evaluated the performance of CelloType on simultaneous segmentation and classification of both cell and non-cell elements in the bone marrow, including small regularly shaped cell types and much larger adipocytes and irregularly shaped trabecular bone fragments. CelloType achieved average AP50 values of 55.4, 44.3, and 58.9 for adipocytes, trabecular bone fragments, and the rest of cell types, respectively (Figure 6a). Consistent with our results with the colorectal cancer CODEX dataset, we observed a strong correlation between the prediction confidence scores and prediction accuracy (Figure 6b). Figure 6c shows two representative examples of predictions by CelloType along with the ground truths. In addition to correctly identifying smaller and regularly shaped cells, CelloType correctly identified most adipocytes and trabecular bone fragments. This result demonstrates CelloType’s efficacy of analyzing challenging tissue images consisting of tightly packed cells and non-cell elements with varying sizes and shapes.

**Figure 6 –.**
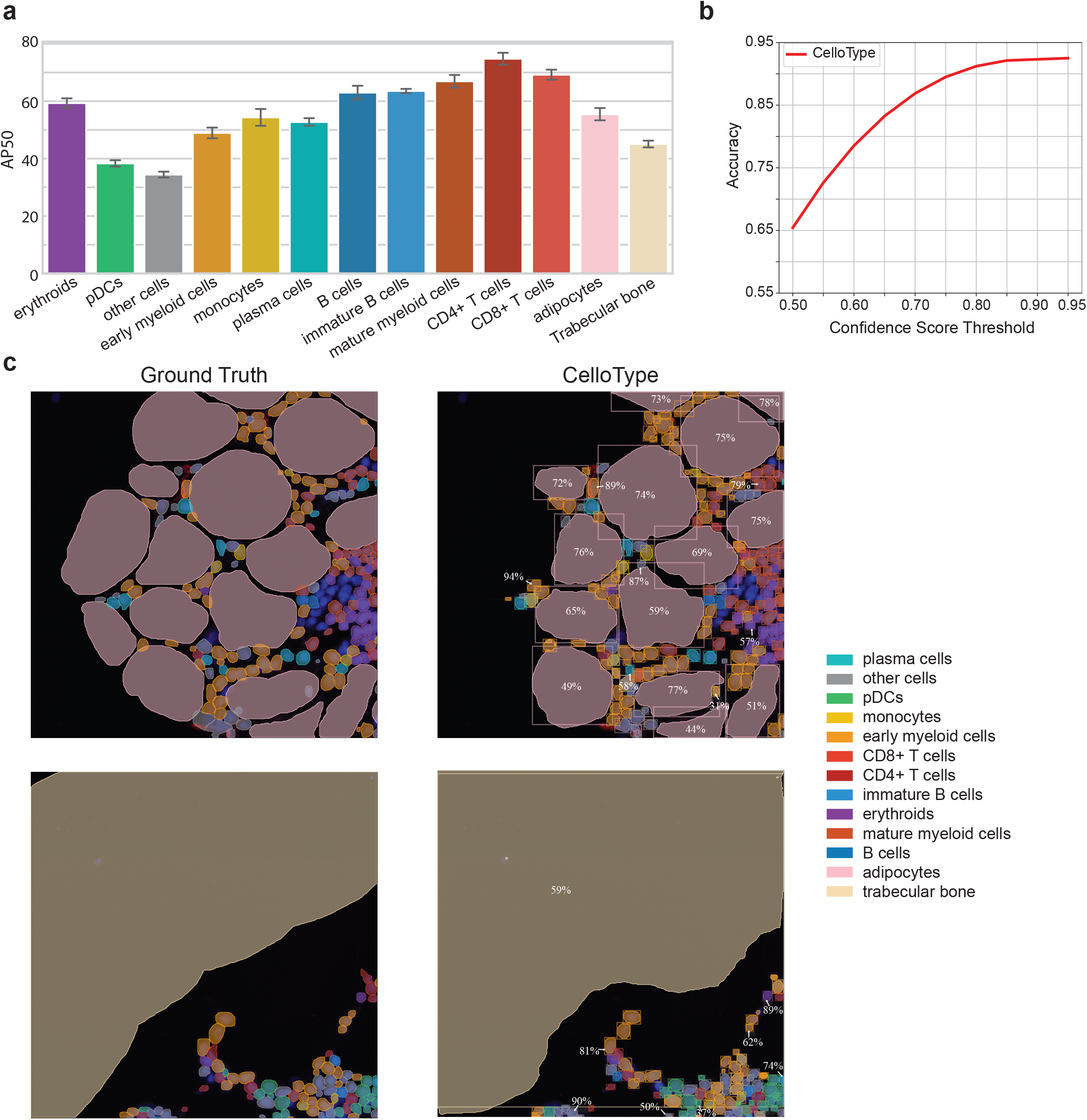
CelloType supports joint multi-scale segmentation and classification. a) Performance evaluation of CelloType stratified by cell and microanatomic structure types. The bar plot shows the mean and 95% confidence interval of AP50 values in 5-fold cross-validation experiments. b) Line plot showing the relationship between classification accuracy and confidence score threshold. c) Representative examples of multi-scale segmentation and classification using human bone marrow CODEX data. The first row of images shows an example of bone marrow area consisting of various types of smaller hematopoietic cells and much larger adipocytes. The second row of images shows an example of bone marrow area consisting of various hematopoietic cell types and microanatomic structure such as trabecula bone fragments. Randomly selected confidence scores for cell classification were displayed next to the predicted instances.

## Discussion

We present CelloType, an end-to-end method for joint segmentation and classification for biomedical microscopy images. Unlike existing methods that treat segmentation and classification as separate tasks, CelloType uses a multi-task learning approach. By leveraging advancements in Transformer-based deep learning techniques, CelloType offers a unified approach to object detection, segmentation, and classification. It starts with Swin Transformer-based feature extraction from an image, followed by the DINO object detection and classification module, which produces latent features and detection boxes that, when combined with the raw image inputs within the MaskDINO module, culminate in refined instance segmentation and classification. The shared encoder in the DINO module extracts latent information that is shared by both tasks, explicitly enhancing the connection between the segmentation and classification tasks, and simultaneously boosting the performance of both tasks. Moreover, the improved object detection accuracy of DINO through deformable attention and contrastive denoising allows the classification task to focus on relevant regions of the image.

It should be noted that this work has the following limitations. First, CelloType requires training for segmentation and classification tasks. In terms of segmentation, there is a rapid growth of training data, exemplified by resources like TissueNet and Cellpose Cyto databases. Models that are pre-trained on these public datasets are readily transferable to new images, provided that they contain nuclear and/or membrane channels. However, for classification, training data is considerably more limited. As a result, pretrained CelloType classification model cannot be readily applied to new images unless there is a substantial overlap of cell/structure types between the training and testing images. To mitigate this need for training data for classification, methodologies such as few-shot learning^28^, self-supervised learning, and contrastive learning^29^ can be incorporated into the CelloType framework. Additionally, with the rapid growth of spatial omics data, it is anticipated that high-quality tissue annotations will also grow quickly.

Consequently, CelloType’s pre-training process can be broadened to include a wider array of datasets, thereby facilitating its application in automated annotation of common tissue types.

Spatial transcriptomics technologies can profile hundreds to thousands of genes at single-cell resolution, yielding a much larger number of features compared to spatial proteomics technologies such as CODEX which typically can only profile fewer than a hundred proteins. This substantial increase in the feature space, coupled with the distinct spatial distribution patterns of RNA transcripts versus proteins, introduces new computational challenges for segmentation and classification. To address the challenge of high dimensionality, a spatially aware dimensionality reduction step^30^ can be integrated into the CelloType framework. To capture the spatial distribution patterns of RNA transcripts, an additional learnable positional embedding step can be introduced in the DINO module. These enhancements could significantly broaden CelloType’s applicability to a wide range of spatial omics data.

## Online Methods

### CelloType

A schematic overview of CelloType is depicted in Fig. 1a. The method consists of three modules: 1) a feature extraction module based on the Transformer deep neural network model to generate multi-scale image features which are used in the DINO and MaskDINO modules; 2) a DINO module for object detection and classification; and 3) a MaskDINO module for segmentation. The resulting latent features and detected bounding boxes are then integrated with the input image in the MaskDINO module to produce instance segmentation results. Both DINO and MaskDINO modules are integrated in a single neural network model for an end-to-end learning.

#### Feature extraction module

Multi-scale image features are generated using the Swin Transformer^17^ deep neural network model. Swin Transformer is a hierarchical version of the original Transformer model that utilizes shifted window operations for efficient self-attention. It can capture both local and global features, outperforming conventional convolutional networks in modeling complex image data with improved computational efficiency. Here we use the Swin-L Transformer model pretrained on the Common Objects in Context (COCO) Instance Segmentation dataset^25^.

#### DINO object detection and classification module

The DINO^14^ deep neural network architecture, standing for "DETR with Improved DeNoising Anchor Boxes", is a novel end-to-end object detection model improving upon the DETR (Detection Transformer) architecture. DINO leverages the strengths of the Transformer architecture to effectively capture spatial relationships, essential for discerning overlapping or adjacent cells. On the other hand, DINO incorporates denoising techniques essential for the precise identification of cells against intricate backgrounds and under-varied imaging conditions. Major components of the DINO architecture in CelloType are described as follows.

1. Query Initialization and Selection: To generate the initial anchor box for detecting objects, the model uses two types of queries: positional queries and content queries. It initializes anchor boxes only based on the positional information of the selected top-K features, while keeping content queries unchanged. These queries provide spatial information of the objects. On the other hand, content queries remain learnable and are used to extract content features from the image. This mixed query selection strategy helps the encoder to use better positional information to pool more comprehensive content features, hence more effectively combines spatial and content information for object detection. This mixed query selection method is formulated as following:

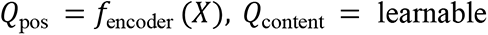

where *Q_pos_* and *Q*_content_ represent positional and content queries, respectively. *Q_pos_* is a n-by-4 matrix and *Q*_content_ is a n-by-embed_dim matrix where n is the number of anchor boxes and embed_dim is the embedding dimension. *X* represents the flattened image features and positional embeddings.

1. Anchor Box Refinement and Contrastive Denoising Training: DINO refines the anchor boxes step-by-step across decoder layers using deformable attention^31^. The conventional attention mechanism examines the whole image whereas the deformable attention selects more important regions of the image and controls the range of self-attention more flexibly, making the computation more efficient. The conventional denoising training technique^32^ involves adding controlled noise to ground truth labels and boxes, formulated as:

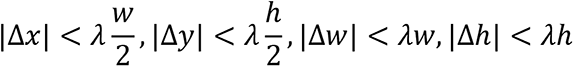

where (*x, y, w, h*) denotes a ground truth bounding box where (*x, y*) is the center coordinates of the box and *λ* and ℎ are the width and height of the box. 𝜆 denotes a hyper-parameter controlling the scale of noise. Contrastive Denoising Training adds both positive and negative samples of the same ground truth, enhancing the model’s ability to distinguish between objects. DINO involves generating two types of queries (positive and negative) with different noise scales *λ*_1_ and *λ*_2_, where *λ*_1_ < *λ*_2_.

1. Classification Head and Confidence Score: For the classification of each bounding box, a linear transformation is applied to the corresponding denoised features. The linear layer outputs a logit vector *Z* = [𝑧_1_, 𝑧_2_, …, 𝑧_𝐾+1_], where K is the number of classes. The vector represents the raw predictions for K classes and the “no object” class. Subsequently, a SoftMax function is employed on the logit vector to compute the class probabilities:

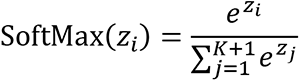

The confidence score for each detected object is taken as the maximum class probability (excluding the “no object” class) outputted by the model. This score represents the model’s confidence in its prediction of the class for the detected object.

#### MaskDINO segmentation module

We use MaskDINO^16^ to predict the segmentation masks using outputs from the feature extractor module and DINO decoder. MaskDINO enhances the DINO architecture by integrating a mask prediction branch. This mask branch utilizes the DINO decoder’s content query embeddings, 𝑞_𝑐_, to perform dot-product operations with pixel embedding maps, derived from both image and latent features at high resolution. These operations result in a set of binary masks, where each segmentation mask, 𝑚, is computed as follows:

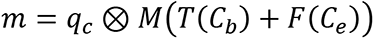

where 𝑞_𝑐_ is the content query embedding, 𝑀 is the segmentation head, 𝑇 is a convolutional layer to map the channel dimension to the Transformer hidden dimension, 𝐶_𝑏_ is the feature map from the feature extractor module, 𝐶_𝑒_ is the latent features from the DINO Transformer encoder, and 𝐹 is an interpolation-based upsampling function to increase the resolution of latent feature and to make the result match the size of the image feature.

Segmentation task, being a pixel-level classification task, offers more detailed information in the initial training stages compared to the region-level object detection task. Therefore, MaskDINO employs the Unified and Enhanced Query Selection technique, which enables the DINO object detection module to leverage the detailed information from the segmentation task early in the training process, enhancing the detection task by providing better-initialized queries for subsequent stages. This cooperative task approach between detection and segmentation results in improved detection performance due to the enhanced box initialization informed by segmentation mask.

During the unified model training, the loss function is calculated by considering three components: segmentation mask, bounding box prediction and class prediction. The composite loss function is expressed as follows:

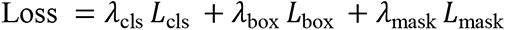

where 𝐿_cls_, 𝐿_box_, 𝐿_mask_ represent classification, bounding box, and segmentation mask losses, respectively, and *λ*_cls_, *λ*_box_, *λ*_mask_ are their corresponding weights.

### Implementation of CelloType for Segmentation tasks

The CelloType software was implemented using the Detectron2 library. Detectron2 is a Facebook AI Research open source library that provides a high-performance, easy-to-use implementation of state-of-the-art object detection algorithms written with PyTorch^33^.

Furthermore, it efficiently manages large datasets and features a flexible architecture that facilitates customization and integration of various image detection or segmentation pipelines.

Dataset was randomly divided into 80% for training, 10% for validation, and 10% for testing. All images and cell/nuclear masks in the training, validation, and testing sets were converted to align with Detectron2’s JSON dictionary schema. For the training dataset, bounding boxes were derived for each cell using the ground truth segmentation masks. The final dictionary encompasses the bounding box, segmentation mask, and raw image for each cell.

For model training, we initialized the DINO and MaskDINO parameters using the weights pretrained on the COCO instance segmentation dataset, as this dataset is extensive and diverse, providing a foundational knowledge for the model. This pretraining helps in better feature extraction and generalization. We used the Adam optimizer with a learning rate of 10^-6^ and a batch size of 8. For every 5 training epochs, the trained model was evaluated on the validation set. The training was terminated when the evaluated AP scores did not improve after 15 epochs. The model with the best AP scores was used for predicting the cell masks.

For evaluation and testing, we set the number of queries to 1,000 which determines the number of boxes and masks generated by the model. In general, this number should exceed the instance count in each image yet remain reasonable to reduce computational cost. Considering the maximum cell count in an image patch in all our datasets does not exceed 1,000, this number was used as the default parameter. Consequently, the model outputs 1,000 instances per image, each comprising a segmentation mask and a corresponding confidence score. For testing, a confidence threshold of 0.3 was used to call predicted instances.

### Implementation of CelloType for classification task

The same training, validation, and testing protocols were used as for the segmentation task. However, during model training for multiplexed images with over three channels, the n_channels hyperparameter within the Swin Transformer was set to match the input images’ dimensionality.

### Running of existing methods

#### Mesmer

Mesmer was run using the pretrained model detailed by the authors in the “Mesmer- Application.ipynb” notebook located in the DeepCell-tf GitHub repository. Key parameter settings included “image_mpp”= 0.5, "compartment” = “whole-cell" for cell segmentation and "nuclear" for nuclear segmentation.

#### Cellpose2

For TissueNet and Cellpose Cyto datasets, Cellpose2 was run using the pretrained model provided by the authors. For colorectal and bone marrow CODEX datasets, we retrained the Cellpose2 model following the procedure described by the authors at https://cellpose.readthedocs.io/en/latest/gui.html#training-your-own-cellpose-model.

#### CellSighter

We trained the CellSighter cell type classification model following the protocol provided by the authors. Key parameters settings included "crop_input_size"=60, "crop_size"=128, “epoch_max”=300 epochs, and “lr”=0.001.

#### Combining Cellpose2 and CellSighter for segmentation and classification

Since there is no existing method for end-to-end joint segmentation and cell type classification, we devised a baseline model combining Cellpose2 and CellSighter, given their reported high performance in the respective tasks. Training of the hybrid model comprised two steps, each optimizing the performance of the individual method. For Cellpose2, CODEX images and corresponding ground-truth cell segmentation masks were used for model training. For CellSighter, the same ground-truth cell segmentation masks along with associated cell type labels were used for training.

During the testing phase, a CODEX image was processed with the trained Cellpose2 model to produce cell segmentation masks, which were subsequently used by the trained CellSighter model for cell type classification. The final results were the combination of the segmentation results of Cellpose2 and cell type classification results of CellSighter.

### Metrics and procedure for evaluating segmentation accuracy

The Average Precision (AP) metric is a widely adopted standard for evaluating the performance of instance segmentation methods in computer vision tasks^34,35^. Specifically, for a given Intersection-over-Union (IoU) threshold, 𝑡, a prediction is considered a true positive if the IoU between the predicted segmentation and the ground truth is greater than 𝑡. The IoU is defined as the ratio of the area of overlap between the predicted segmentation mask and the ground truth mask. The AP is calculated at IoU values from 0.50 to 0.9 with a step size of 0.05. The final AP is the average of the AP values at these different IoU thresholds. This gives a more comprehensive understanding of a model’s performance, from relatively lenient (IoU=0.50) to stricter overlaps (IoU=0.9).

In the context of multiple classes, mAP is computed by taking the mean of the AP values calculated for each individual class. Specifically, if the task only has one class, such as cell segmentation or nuclear segmentation, the mAP would be the average precision across all the IoU we evaluated. This gives an overall sense of the method’s performance across the various classes in the dataset, rather than focusing on its efficacy in detecting a single class.

To evaluate segmentation performance using the AP metric, we used the Common Objects in Context (COCO) evaluation package, a widely used, standardized benchmarking tool in the field of instance segmentation. Segmentation results were first converted into the COCO format before the AP metric was computed using the package. To eliminate redundant detections and ensure that each object is uniquely identified, the package implements the Non-Maximum Suppression (NMS) procedure. NMS selectively filters out overlapping bounding boxes, retaining only the box with the highest confidence score while discarding others with substantial overlap, as determined by the IoU threshold. Since methods such as Mesmer, Cellpose2, and CelloType do not generate confidence score for the predicted segmentation masks, we arbitrarily assigned the confidence score to be 1. For the CelloType variant that outputs the confidence score (CelloType_C), we used the actual confidence scores computed by the method when applying the NMS procedure.

## Datasets

### TissueNet dataset

The TissueNet dataset^4^ consists of 2,601 training and 1,249 test multiplexed images collected using multiple imaging platforms and tissue types. Imaging platforms include CODEX, CycIF, IMC, MIBI, MxIF and Vectra. Tissue types include breast, gastrointestinal, immune cells, lung, pancreas, and skin. Although many images have dozens of protein markers, all images contain at least two channels necessary for cell/nucleus segmentation: a cell membrane channel and a nuclear channel. Each image contains a manual segmentation of cells and/or nuclei. Each training and test image has a dimension of 512 × 512 pixels and 256 × 256 pixels, respectively.

### Cellpose Cyto dataset

The Cyto dataset^6^ consists of images from a variety of sources, including: 1) Cells (Cell Image Library) set: 100 fluorescent images of cultured neurons with both cytoplasmic and nuclear stains obtained from the Cell Image Library database (http://www.cellimagelibrary.org); 2) Cells (Fluorescent) set: 216 fluorescent images of cells visualized with cytoplasmic markers. This set contains images from BBBC020, BBBC007v1, mouse cortical and hippocampal cells expressing GCaMP6 imaged using a two-photon microscope, confocal images of mouse cortical neurons, and the rest were obtained through Google image search; 3) Cells (non-fluorecent) set: 50 brightfield microscopy images from OMERO and Google image search; 4) Cells (Membrane) set: 58 fluorescent images of cells with membrane maker, 40 of which were from the Micro-Net image set and the rest were obtained through Google image search; 5) Other microscopy set: 86 images of other types of microscopy that contain either non-cells or cells with atypical appearances. These images were obtained through Google image search; 6) Non-microscopy set: 98 images of non-microscopy images obtained through Google search of repeating objects including images of fruits, vegetables, artificial materials, fish and reptile scales, starfish, jellyfish, sea urchins, rocks, seashells, etc. All images in the dataset were manually segmented by a human operator.

### Colorectal cancer CODEX dataset

This dataset contains CODEX images of 140 human colorectal samples stained with a 56 fluorescent antibodies and 2 nuclear stains^26^. Cells were segmented using Mesmer. Cell types were annotated by the authors using a combination of iterative clustering and manual examination of marker expression profiles and morphology. For each tissue image in the dataset, image patches of 512 x 512 pixels were generated.

### Bone marrow CODEX dataset

This dataset^27^ contains CODEX images of 12 human bone marrow samples stained with 54 fluorescent antibodies and one nuclear stain. Hematopoietic cell types were annotated by the authors using a combination of iterative clustering and manual examination of marker expression profiles and morphology. Adipocytes and trabecular bone fragments were manually annotated by the authors.

## Code availability

CelloType is available at: https://github.com/tanlabcode/CelloType.

**Supplemental Figure 1 – Distribution of cell types in the colorectal cancer CODEX dataset.**

**Supplemental Figure 2 – Distribution of cell types in the human bone marrow CODEX dataset.**

## Acknowledgments

The authors thank the Children’s Hospital of Philadelphia Research Information Services for providing computing support. This work was supported by the National Cancer Institute (NCI) Human Tumor Atlas Network grant under award #U2C CA233285 (K.T.) and the National Institutes of Health (NIH) Human Biomolecular Atlas Program grant under award #U54 HL165442 (K.T.).

## Author Contributions

M.P. and K.T. conceived and designed the study. M.P. implemented the CelloType algorithm and wrote the software, M.P. and T.R. performed data analysis, X.W. and K.T. supervised the overall study. M.P. and K.T. wrote the manuscript with input from all authors.

